# Light-Controlled Synthetic Communication Networks via Paired Connexon Nanopores

**DOI:** 10.1101/2025.04.11.648033

**Authors:** Ahmed Z. Sihorwala, Alexander J. Lin, Isabela Ramirez-Velez, Siqi Yang, Shwetadwip Chowdhury, Jeanne C. Stachowiak, Brian Belardi

## Abstract

Living cells employ dynamic networks for intercellular communication and cooperation, leading to tissue-wide activity. One emerging challenge in the field of bottom-up synthetic biology is emulating such sophisticated behaviors in liposome-based synthetic cells (SCs). Fabricating communication networks in lipid bilayer-based SCs remains a challenge as signaling molecules must transit through two consecutive membranes to transfer information between different SCs. Here, we address this obstacle by engineering connexin channels that directly connect the lumens of adhering SC membranes. We focus on orthogonal channel-forming connexins, namely connexin 43 and connexin 32, and re-design their channel activity to be UV- and near IR-responsive, respectively. By combining engineered connexins into a single SC assembly, we demonstrate orthogonal transfer of reactive signaling molecules between SCs, giving rise to unique reaction products and network states in a wavelength-dependent manner – an important step toward synthetic communication networks.

## Introduction

The movement of electrons, ions, and molecules underlie everyday networks capable of long-range communication. In living systems, the transfer of molecular signals – metabolites, hormones, and growth factors – between cells forms one basis of network communication that is critical for collective and coordinated action,^1^ for example during quorum sensing^2^ and tissue differentiation and morphogenesis.^3^ To ensure robustness and fidelity in an environment of competing chemical signals, living cells exhibit tight control over when, where, and how molecular signals are transferred and interpreted.^4,5^ For instance, sender cells release signals at precise times and locations, and through signal transduction mechanisms or expression of transporters or channels, receiver cells process those released molecular signals. In higher animals with a multitude of cell types, signal transfer requires specificity, such that multiple communication networks can act simultaneously between subsets of cells across tissues and organs without interference.^6^ Emulating living communication networks in synthetic systems would lead to new classes of materials that transfer and transform an array of molecular information dynamically, critical features for regenerative medicine, advanced sensors, and computing.

One synthetic platform that offers distinct advantages for material construction is synthetic cells.^7^ Here, we define synthetic cells (SCs) as micron-sized lipid vesicles that mimic cellular form, properties, and functions. Their programmable nature, compartmentalization, membrane functionality, and bio- and aqueous solvent-compatibility have made SCs attractive building blocks for a variety of applications.^8–11^ These properties have also been leveraged for signal release.^12–20^ By virtue of their lipid bilayer boundaries, compartmentalization of a wide range of hydrophilic molecules in SCs is routine,^21^ which has subsequently enabled release of stored signals upon expression of membrane pores, such as the bacterial α-hemolysin.^22–25^ However, these same SC properties make high fidelity, directed transfer far more challenging. Signals are often released into the extracellular milieu, where they indiscriminately reach nearby SCs in a concentration-dependent manner, limiting both specific transfer and fidelity as some signal is lost to the surrounding medium, the latter of which reduces signal amplitude. User-defined, direct transfer of signals from one SC to another would avoid these drawbacks, opening up the possibility of orthogonal communication between SCs. Such a capability would enable multiple SC assembly states, yet remains elusive to date.

A conduit structure that passes through two lipid bilayers would transform SCs into a network capable of direct molecular signal transfer.^26,27^ Taking inspiration from cellular gap junctions, we report the first example of a user-controlled protein conduit system between SCs. Gap junctions depend on a family of proteins known as connexins that self-assemble into hexameric nanopores, so-called connexons, in cell membranes.^28–30^ Connexon nanopores can couple across cells, forming full channels that exchange cytoplasmic contents between connected neighbors.^31,32^ Here, we demonstrate full connexin channel activity for both connexin 43 (Cx43) and connexin 32 (Cx32) between SCs and find that the presence of adhesive co-receptor proteins is necessary for efficient signal transfer. To impart user-control over signal transfer, we further engineer Cx43- and Cx32-based connexons, such that their nanopore assembly is caged by a bulky domain. To activate connexon assembly, we take advantage of protease cleavage of the bulky domain, which we couple to either UV or near-IR illumination. With this system, channel formation of either Cx43 or Cx32 channels across adhered SCs can be selectively triggered with different wavelengths of light. We showcase how our SC network can be used to transfer specific reactive molecules between SCs, in turn producing unique reaction products and SC assembly states in a wavelength-dependent fashion, akin to signaling-based differentiation in a tissue. Our work constitutes an important step towards constructing synthetic communication networks with high fidelity, specificity, and user-defined control that approach the features of living tissue.

## Results and Discussion

### Adhesive contacts enhance connexin channel formation between synthetic cells

Connexin channels form in living cells when connexon extracellular loops bind across cells in a trans fashion.^33^ Here, we sought to assemble connexin channels between SCs as a means to facilitate inter-SC signal transfer. To do this, we created two SC populations – one encapsulating a soluble dye (sender SCs) and another lacking an intracellular dye (receiver SCs). The dye acts as a molecular signal to monitor transfer between SCs. To prevent inter-SC transfer via dye leakage from unpaired connexons, we added La^3+^, a known unpaired connexon blocker,^34,35^ to the SC outer solution (Supplementary Fig. 1). Upon expression of Cx43 in both sender and receiver SCs using a cell-free protein expression system (PURE system), we observed a modest increase in dye transfer to receiver SCs that express Cx43 (+pCx43, 20% dye transfer) in comparison to those that do not (-pCx43, 12% dye transfer) (Fig. 1b, c). We attributed minimal channel-based transfer to inadequate contact formation between sender and receiver SCs as connexons’ extracellular loops undergo fast dissociation and slow association.^36^ We, therefore, reasoned that an additional adhesive complex that can form stable membrane contacts between SCs may increase the pairing probability of connexons.

**Figure 1.**
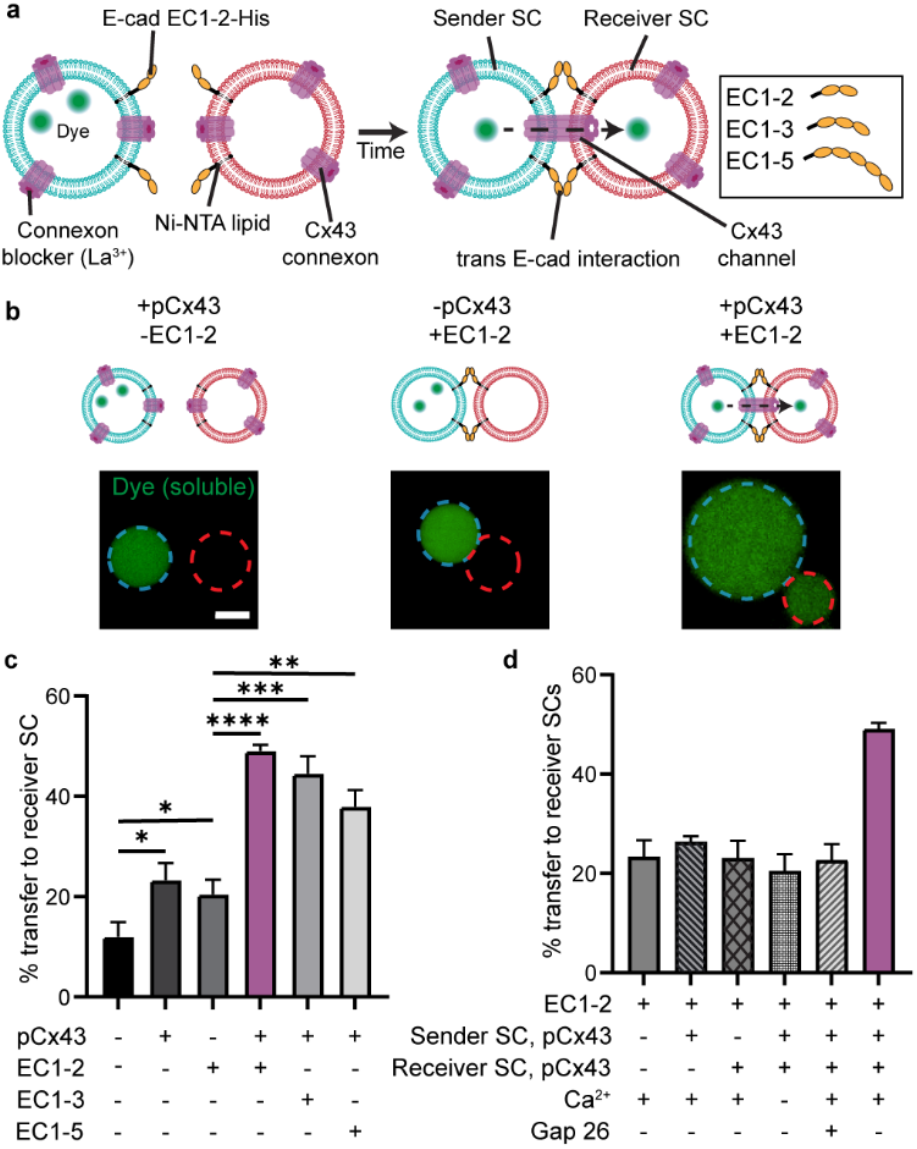
Adhesion-mediated assembly of Cx43 channels between SCs. **a**, Scheme of Cx43 connexin channel assembly between sender (cyan) and receiver (red) SCs. SC membranes incorporating DGS-NTA(Ni) are functionalized with different size variants of His-tagged E-cadherin ectodomain (EC1-2, EC1-3, or EC1-5). Cx43 channels between sender and receiver SCs lead to transfer of dye (green, Alexa Fluor 647) between SCs. La^3+^ blocks any unpaired connexon, ensuring dye transfer is connexin channel-dependent. **b**, Fluorescent micrographs of sender and receiver SCs demonstrate that dye transfer is dependent on EC1-2-mediated adhesion between Cx43-expressing SCs. **c**, Quantification of dye transfer to receiver SCs in the presence of different E-cadherin variants (EC1-2, EC1-3, or EC1-5) reveals maximum transfer occurs between Cx43-expressing SCs that are adhered by EC1-2 (highlighted in purple). Error bars represent the s.d. of 3 independent trials, at least 35 receivers were analyzed per trial. **d**, Quantification of dye transfer between EC1-2-tethered SCs in the presence or absence of Cx43 and with and without a Cx43-channel inhibitor (Gap26) or extra-SC Ca^2+^, the latter of which is necessary for EC1-2-mediated adhesions. Without Cx43 or Ca^2+^ or in the presence of Gap26, dye transfer was abolished. Error bars represent the s.d. of 3 independent trials, at least 35 receivers were analyzed per trial. Scale bar: 5 µm. Asterisks represent statistically significant differences (two-tailed unpaired t test, *p < 0.05, **p < 0.01, ***p < 0.001, ****p < 0.0001, n.s. p > 0.05).

To test this hypothesis, we functionalized sender and receiver SC membranes with the extracellular domains of the adherens junction protein, E-cadherin (E-cad). E-cad has five ecto-domains with only the first two domains associating in trans to adhere neighboring epithelial cells.^37^ We recombinantly purified a hexahistidine-tagged (His-tag) version of E-cad with its adhesive domains (EC1-2) to induce SC adhesion. To tether EC1-2 to sender and receiver SC surfaces, we doped SC membranes with 5 mol% DGS-NTA(Ni) lipid. Following mixing and a pelleting step, robust contacts formed between sender and receiver SCs (Supplementary Fig. 2). In the presence of EC1-2, significant dye transfer (50%) was observed between Cx43-expressing sender and receiver SCs (+pCx43) compared to SCs that did not express Cx43 (-pCx43, 23%) (Fig.1a-c). Our data with EC1-2 suggests that adhesion between SCs increases the probability of connexon pairing by bringing SC membranes in close proximity, leading to robust signal transfer.^38^ To probe this effect further, we decorated sender and receiver SCs with longer versions of His-tagged E-cad (EC1-3 and EC1-5). We observed a reduction in dye transfer to receiver SCs with the longer variants of E-cad (Fig. 1c) compared to EC1-2 despite similar binding energies at SC interfaces (Supplementary Fig. 2). Energetically costly membrane bending may explain the decrease in transfer as the E-cad construct length increases. Full Cx43 channel formation leads to an intermembrane distance of ∼3.5 nm,^39^ whereas full-length E-cad adhesions position membranes ∼37 nm apart.^37^ Using the short EC1-2 variant would reduce this distance to ∼25 nm,^40^ leading to less membrane bending. Since the short EC1-2 variant showed the highest extent of inter-SC transfer, we used it for all subsequent experiments involving connexin channel assembly.

Next, we optimized the concentration of EC1-2 that drove maximum dye transfer (50 nM) (Supplementary Fig. 3) and ensured that transfer was contingent on both populations of SCs expressing Cx43 (Fig. 1d). As well, we verified that adding a Cx43-extracellular loop mimetic peptide, Gap26,^41^ a known channel inhibitor, to the extra-SC solution, reduced dye transfer. Finally, EC1-2’s trans interactions require Ca^2+^.^42^ Upon removing Ca^2+^ from the extra-SC solution, only background transfer occurred, confirming that enhanced transfer was indeed Cx43 channel-dependent and due to EC1-2-mediated contacts between SCs. Together, our data show that connexin channels can form between micron-sized SCs and that transfer can be enhanced by engineering SCs with short adhesive co-receptors.

### Pairing-incompatible connexons form orthogonal channels and lead to non-mixing transfer paths

Having found that Cx43 forms channels across adhered SCs, we next examined whether we could extend this approach to another connexin isoform, especially one that may form orthogonal channels to Cx43, potentially lending specificity to SC communication. Towards this end, we focused on assembling connexin 32 (Cx32) channels between SCs, which are thought to be docking-incompatible with Cx43 based on data from live cell experiments.^43,44^ By encapsulating the PURE system along with a Cx32-GFP plasmid (pCx32) inside SCs, we were first able to assemble functional Cx32 connexons that released dye to the surrounding solution (Supplementary Fig. 4). Using EC1-2 to drive adhesion between sender and receiver SCs, we then found evidence of adhesion-mediated dye transfer across Cx32 channels (Fig. 2a, Supplementary Fig. 5), albeit to a lower extent than for Cx43 channels (Fig. 1). This observation is in agreement with previous work in cells that has shown lower permeability and conductance of Cx32-Cx32 channels compared to their Cx43-Cx43 channel counterparts.^45^ Moreover, expression of Cx32 was lower in SCs compared to Cx43 (Supplementary Fig. 6), which may also contribute to the difference in dye transfer across connexin channels. Notably, when we paired sender SCs expressing Cx43 with receiver SCs expressing Cx32, dye transfer was abolished (Fig. 2c), in line with the cell-based work,^43,44^ and most likely due to differences in key hydrogen bonding residues that reside in the second extracellular loops (ECL2) of Cx43 (H194) and Cx32 (N175) (Fig. 2b).^33^ To probe this incompatibility further, we designed a three SC population experiment, wherein a sender SC that expresses either Cx43 or Cx32 and encapsulates a dye is combined with both a Cx43-expressing receiver SC and a Cx32-expressing receiver SC. After mixing and adhering the three SC populations to one another through EC1-2, we found significant dye transfer only between sender SC and receiver SC that expressed the same connexin isoform (Fig. 2d). Different connexons with incompatible pairing, as we have shown for Cx43 and Cx32, open up the possibility of defining specific communication pathways in SC assemblies.

**Figure 2.**
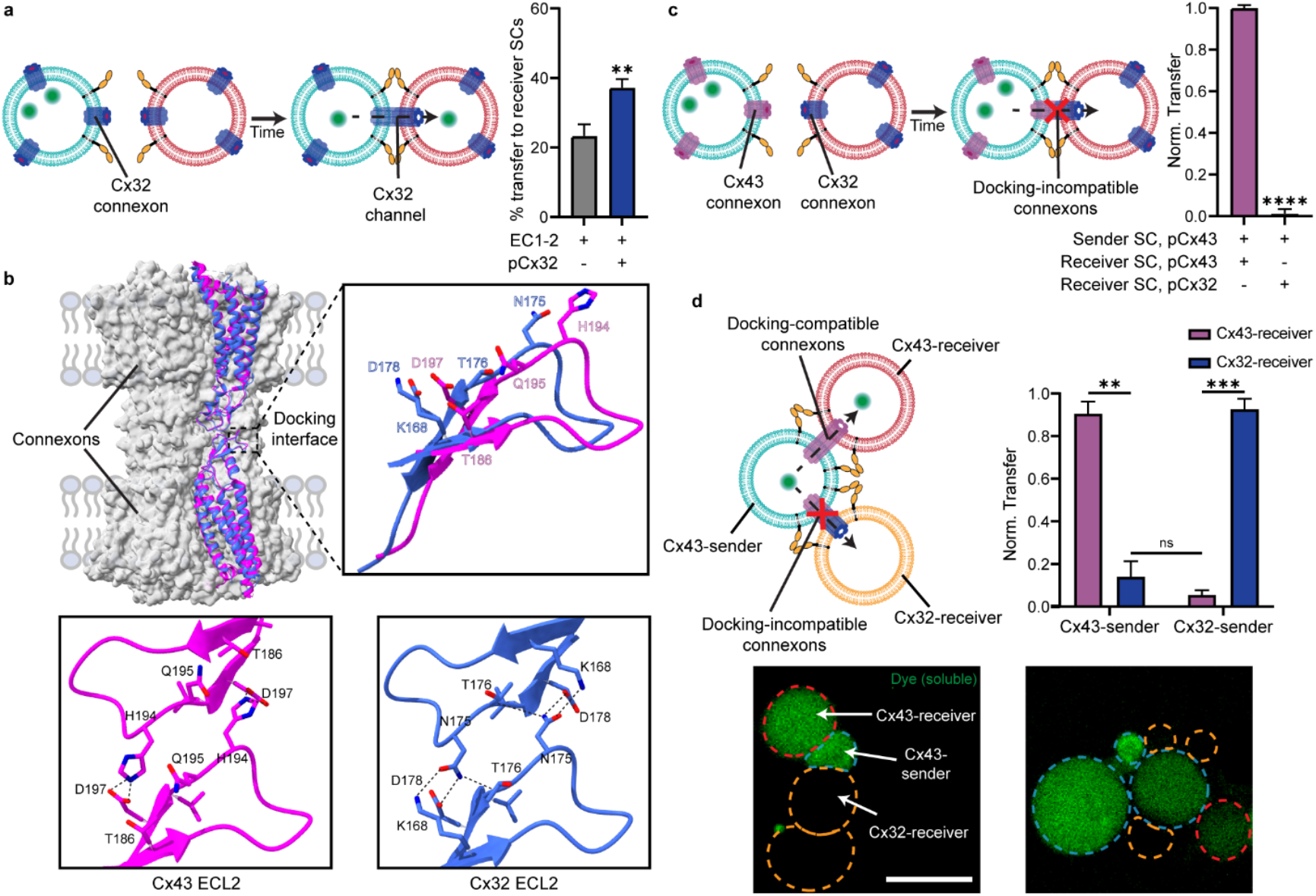
Pairing of orthogonal connexin channels between SCs. **a**, Scheme of adhering SCs with Cx32 channels (left). Quantification of dye transfer to receiver SCs across Cx32 channels (right). Error bars represent the s.d. of 3 independent trials, at least 35 receivers were analyzed per trial. **b**, Superposition of cryo-EM structures of Cx43 and Cx32 channels (top, left). ECL2 regions of Cx43 (pink) and Cx32 (blue) represented as ribbon diagrams, along with key residues involved in inter-connexon interactions of Cx43 and Cx32 channels shown as sticks and labeled (top, right). Differences in the amino acid residues at the respective ECL2 docking interfaces of Cx43 (pink) and Cx32 (blue) (bottom) result in their docking incompatibility. **c**, Scheme of incompatible Cx43 and Cx32 connexons, which would lead to a lack of dye transfer between SCs (left). Normalized dye transfer from Cx43-expressing sender SCs to receiver SCs expressing either Cx43 or Cx32 (right). Error bars represent the s.d. of 3 independent trials, at least 50 receivers were analyzed per trial. **d**, Scheme of selective dye (Alexa Fluor 488, green) transfer between senders (cyan) and receivers (red and orange) in a three SC population based on the pairing compatibility of connexons (top left). Fluorescent micrographs of Cx43-senders (cyan) adhered to Cx43-(red) and Cx32-receivers (orange) show preferential dye transfer to Cx43-receivers (bottom). Normalized dye transfer between Cx43- and Cx32-senders to receivers expressing either Cx43 or Cx32 (top right). Error bars represent the s.d. of 3 independent trials, at least 75 receivers were analyzed per trial. Scale bar: 5 µm. Asterisks represent statistically significant differences (two-tailed unpaired t test, **p < 0.01, ***p < 0.001, ****p < 0.0001, n.s. p > 0.05).

### Light-activated assembly of Cx43 channels between synthetic cells

Achieving direct signal transfer between SCs for user-defined communication would require a means of control over the timing of transfer. We decided to make use of our previously engineered version of Cx43, whose activity was dependent on TEV protease activity.^19^ Engineered Cx43 was designed with a bulky N-terminal domain followed by a TEV protease recognition site (TEVrecCx43), which we previously showed blocks assembly of a functional connexon nanopore.^19^ We also showed previously that by encapsulating TEV protease in UV-responsive liposomes within SCs that connexon pore activity could be triggered with UV light (UV-Cx43).^19^ The sequence of events is as follows: UV illumination causes liposome rupture in SCs, releasing TEV protease. TEV protease then acts on TEVrecCx43, digesting the N-terminal bulky domain, which leads to nanopore assembly and activity. Here, we asked whether we could extend this system to assemble functional Cx43 channels across SCs in response to UV light. To test this concept, we created UV-Cx43 sender and UV-Cx43 receiver SCs. In the absence of light, we found negligible dye transfer between sender and receiver SCs that were adhered with EC1-2, indicative of non-functional connexin channels. However, following illumination with UV light, a significant increase (85-fold) in dye transfer was observed (Fig. 3a). The amount of transfer was indistinguishable from TEVrecCx43-expressing sender and receiver SCs that encapsulated soluble TEV, suggesting efficient light-mediated activation of Cx43 channel activity.

**Figure 3.**
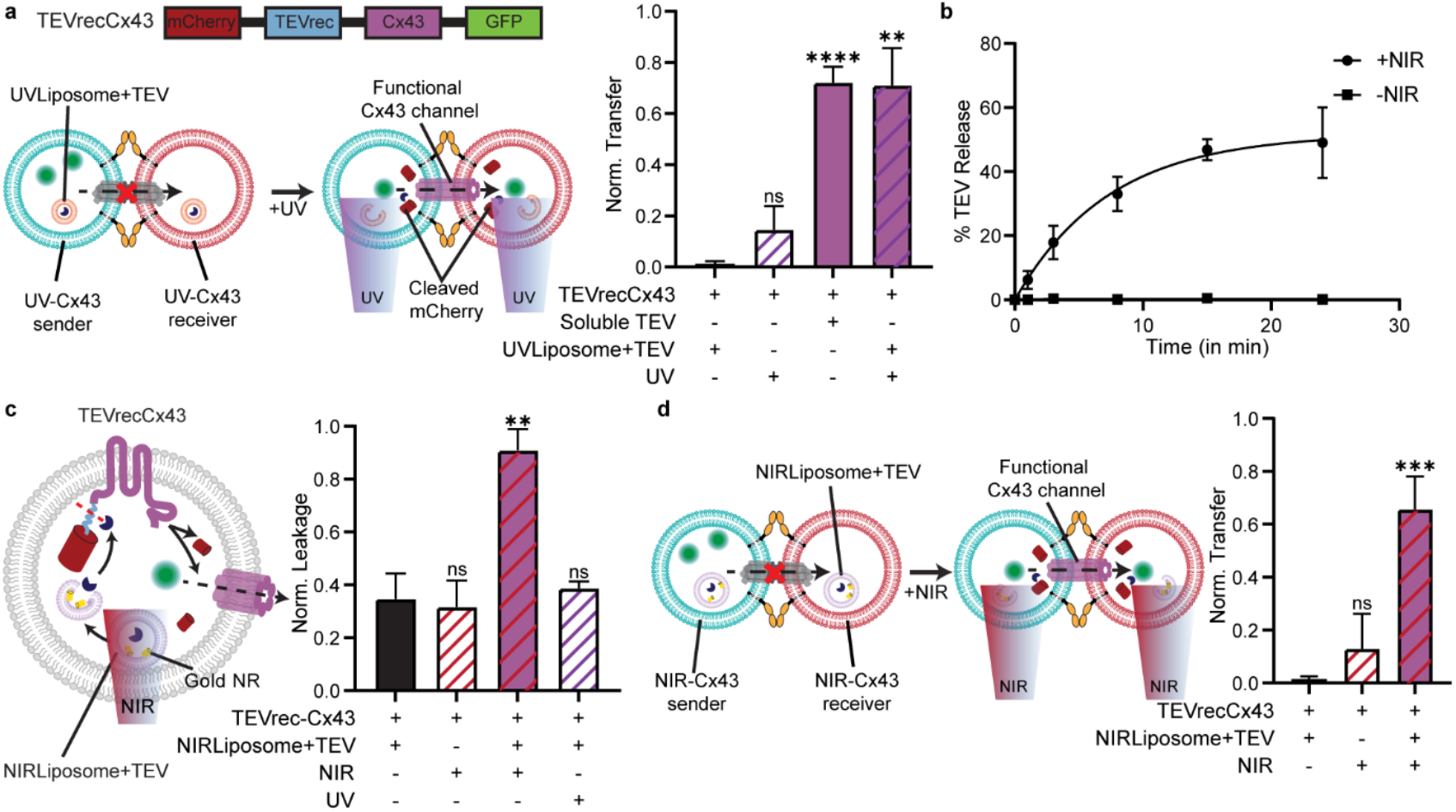
Light-activated assembly of Cx43 channels between SCs. **a**, Primary sequence of TEVrecCx43 (top, left). Scheme of UV-dependent assembly of functional Cx43 channels between SCs (bottom, left). UV-responsive liposomes (UVLiposomes+TEV}) and Alexa Fluor 647 were co-encapsulated in SCs expressing the TEVrecCx43 variant. UV illumination would result in TEV release from liposomes and cleavage of mCherry, leading to dye transfer. Normalized dye transfer to UV-Cx43 receivers in the presence or absence of UV-responsive liposomes or UV illumination (right). Error bars represent the s.d. of 3 independent trials, at least 45 receivers were analyzed per trial. **b**, Percentage of TEV release from NIR-responsive liposomes as a function of illumination time (t_1/2_ = 15 min). **c**, Scheme of NIR-dependent assembly of functional Cx43 connexon in SCs (left). NIR-responsive liposomes containing gold nanorods (NIRLiposomes+TEV) were co-encapsulated with Alexa Fluor 647 in SCs expressing the TEVrecCx43 variant. Upon NIR illumination, TEV release would lead to mCherry cleavage and subsequent dye leakage. Normalized dye leakage in SCs expressing TEVrecCx43 in the presence and absence of NIR-responsive liposomes and NIR illumination (right). Error bars represent the s.d. of 3 independent trials, at least 30 SCs were analyzed per trial. **d**, Scheme of NIR-dependent dye transfer across SCs expressing TEVrecCx43 (left). Normalized dye transfer between NIR-Cx43 senders and receivers in the presence or absence of NIR-responsive liposomes and NIR illumination (right). Error bars represent the s.d. of 3 independent trials, at least 35 receivers were analyzed per trial. Asterisks represent statistically significant differences (two-tailed unpaired t test, **p < 0.01, ***p < 0.001, ****p < 0.0001, n.s. p > 0.05).

With the ability to control connexin channel activity with UV light, we next focused on developing additional wavelength options for activation that would enable the use of multiple connexin isoforms in parallel. A near IR (NIR, λ = 850 nm)-activatable system would be advantageous as its red shift provides an orthogonal mode of activation to the UV (λ = 254 nm)-activated system. For this purpose, we synthesized NIR-responsive liposomes by encapsulating gold nanorods (gold NR) inside 200 nm-sized liposomal membranes composed of 1,2-dipalmitoyl-sn-glycero-3-phosphocholine (DPPC) (Supplementary Fig. 7b) and included TEV protease. In these liposomes, the gold NRs act as photothermal agents to convert NIR light into heat, rupturing DPPC-based liposomes and leading to TEV release in the process.^46^ We found minimal TEV release from the NIR-responsive liposomes in the absence of NIR (Fig. 3b). However, upon NIR illumination, rapid TEV activity from NIR-responsive liposomes occurred, with maximum release and activity at 15 minutes. Building on these results, we next tested whether we could use NIR-responsive liposomes to trigger connexon activity in SCs. Similar to the UV system, we encapsulated NIR-responsive liposomes containing TEV protease inside SCs expressing TEVrecCx43 and observed connexon-based release only upon NIR illumination (2.6-fold over no illumination condition) (Fig. 3c). As anticipated, these SCs were unresponsive to UV light, demonstrating activation orthogonality, and dependent on the presence of gold NRs within the liposomes for connexon activation (Supplementary Fig. 8). We then turned our attention to channel-based transfer. After forming contacts with EC1-2, significant NIR-mediated transfer was observed (70-fold greater than the no illumination condition) (Fig. 3d). We confirmed that both NIR- and UV-dependent SC transfer were unresponsive to the other wavelength (Supplementary Fig. 9), which demonstrates wavelength-selective activation of connexin channels.

### NIR light-activated assembly of TVMV protease-dependent Cx32 channels

Having established that Cx43 and Cx32 do not pair with each other across SCs (see above), we envisioned a synthetic communication network, wherein incompatible connexins mediate separate, high fidelity communication pathways. To control pathway formation independently, connexins that respond to different stimuli, such as distinct wavelengths of light, would be advantageous. Towards this end, we sought to engineer Cx32, such that its assembly was protease-dependent and orthogonal to that of TEVrecCx43. To do this, we examined the cryo-EM structures of Cx32 connexons (Fig. 4a).^47^ Similar to Cx43, the Cx32 N-terminus forms key contacts in the inner lining of the pore structure, which may be leveraged for blocking pore activity by adding N-terminal bulk. To test this idea, we cloned a TVMV protease recognition site between an N-terminal bulky mCherry and Cx32 (TVMVrecCx32). TVMV is a viral protease that possesses similar activity to TEV protease but acts specifically on a distinct peptide sequence, resulting in minimal cross-reactivity between the two proteases (Supplementary Fig. 10a).^48^ We found that TVMVrecCx32-expressing SCs showed significantly reduced dye leakage compared to SCs expressing wildtype Cx32, suggesting that pore formation is largely blocked prior to cleavage of bulky mCherry. When TVMV protease was added to SCs, cleavage of the mCherry domain by TVMV protease led to recovery of connexon activity (Fig. 4b). Motivated by this finding, we next sought to place Cx32 activation under control of NIR light. To do this, we generated NIR-responsive liposomes loaded with TVMV. We then encapsulated NIR-responsive liposomes inside TVMVrecCx32-expressing SCs (NIR-Cx32) and quantified connexon activity in the presence or absence of 15 minutes of NIR light. We found that Cx32-based connexon activity in NIR-Cx32 SCs increased by 3.5-fold following NIR illumination (Fig. 4b). We also confirmed that NIR-Cx32 did not activate in the presence of TEV and that TEVrecCx43 remained non-functional in the presence of TVMV, confirming orthogonality of the two systems (Supplementary Fig. 10b). To examine SC transfer, we assembled both receiver and sender SCs with the NIR-Cx32 system and indeed found that transfer took place only after NIR illumination (∼35-fold over no illumination case) (Fig. 4c). Collectively, these results suggest that Cx32’s transfer activity can be engineered to be light sensitive and that its activation can be manipulated in a manner orthogonal to Cx43’s activation.

**Figure 4.**
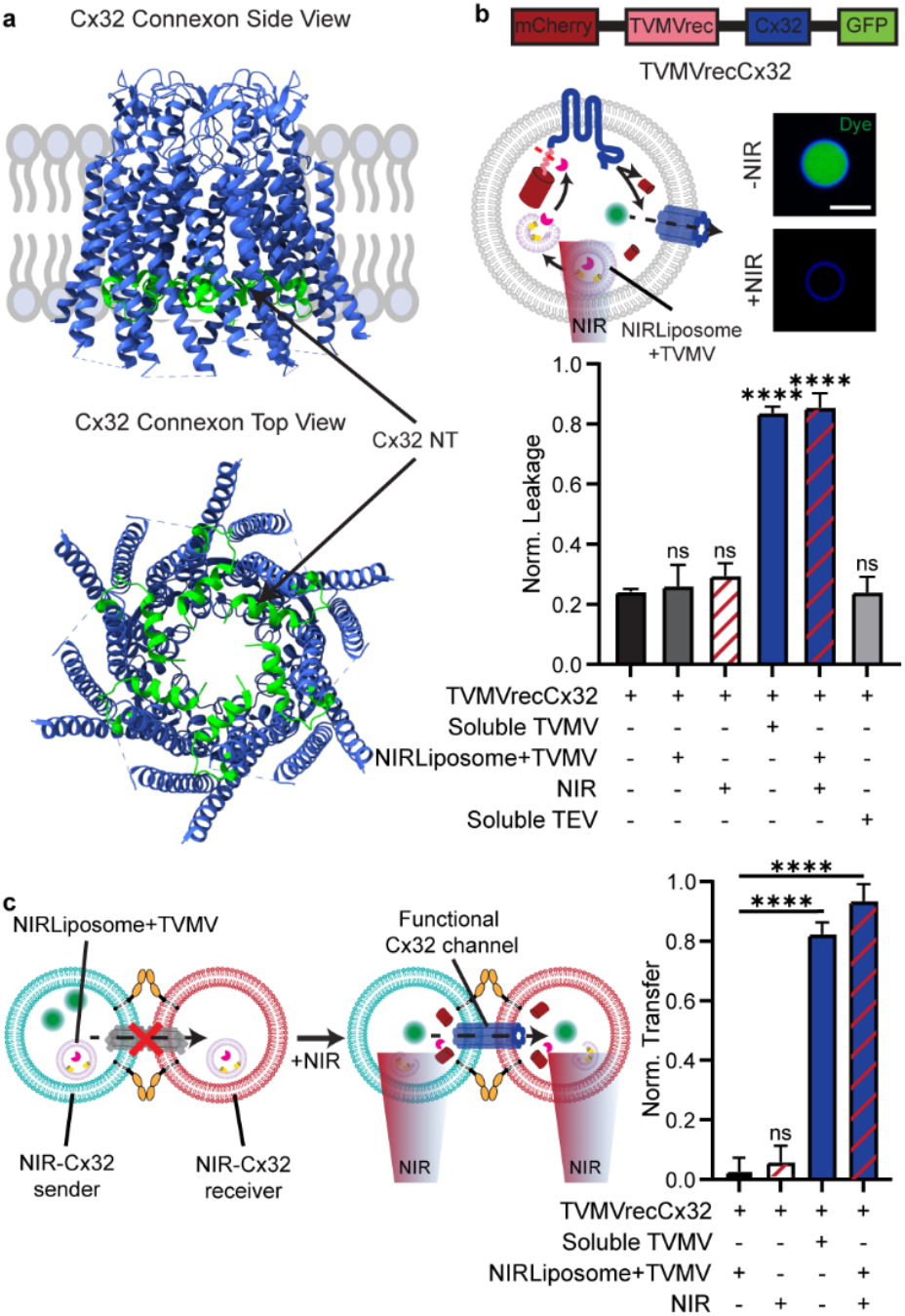
NIR-activated assembly of Cx32 channels between SCs. **a**, Side and top views of Cx32 connexon, PDB 7ZXN. The N-terminus of Cx32 monomers is highlighted in green and faces the interior of the pore. **b**, Primary sequence of TVMVrecCx32 (top). Scheme of NIR-dependent assembly of functional Cx32 connexons in SCs (middle, left). NIR-responsive liposomes with TVMV protease (NIRLiposomes+TVMV) and Alexa Fluor 647 were co-encapsulated in SCs expressing a protease-sensitive variant of Cx32 (TVMVrecCx32). TVMV released from liposomes upon NIR illumination would lead to cleavage of mCherry and subsequent dye leakage from SCs. Fluorescence micrographs showing dye leakage from these NIR-Cx32 SCs following NIR illumination. Normalized dye leakage in SCs expressing TVMVrecCx32 in the presence and absence of NIR-responsive liposomes and NIR illumination (bottom). Error bars represent the s.d. of 3 independent trials, at least 30 SCs were analyzed per trial. **c**, Scheme of NIR-dependent dye transfer across SCs expressing TVMVrecCx32 (left). Normalized dye transfer between NIR-Cx32 senders and receivers in the presence and absence of NIR-responsive liposomes and NIR illumination (right). Error bars represent the s.d. of 3 independent trials, at least 35 receivers were analyzed per trial. Scale bar: 10 µm. Asterisks represent statistically significant differences (two-tailed unpaired t test, ***p < 0.001, ****p < 0.0001, n.s. p > 0.05).

### User-controlled orthogonal communication networks between synthetic cells

Given that the UV-Cx43 and NIR-Cx32 systems are dependent on different proteases for activity, we reasoned that both engineered connexins can be controlled orthogonally in a single SC with separate wavelengths of light. To examine this possibility, we created SCs that contain both UV-Cx43 and NIR-Cx32 systems (DualCx). We then adhered DualCx sender SCs with either Cx43- or Cx32-expressing receiver SCs and, in both cases, found minimal dye transfer (Supplementary Fig. 11), implying an initial “off” state for the DualCx sender. DualCx sender SCs were then adhered with both UV-Cx43 and NIR-Cx32 receiver SCs, constructing a three population SC assembly. As expected, in the absence of light we saw minimal dye transfer between the DualCx sender SC and either receiver SC. Upon UV illumination, however, we observed selective transfer to UV-Cx43 receivers (25-fold over no illumination case) over NIR-Cx32 receivers. Similarly, NIR illumination resulted in predominant transfer to NIR-Cx32 receivers (15-fold over no illumination condition) over UV-Cx43 receivers (Fig. 5). Thus, by using the appropriate wavelength of light, our system allows the user to dictate transfer preferences between SCs, in turn defining independent signaling paths in an SC assembly.

**Figure 5.**
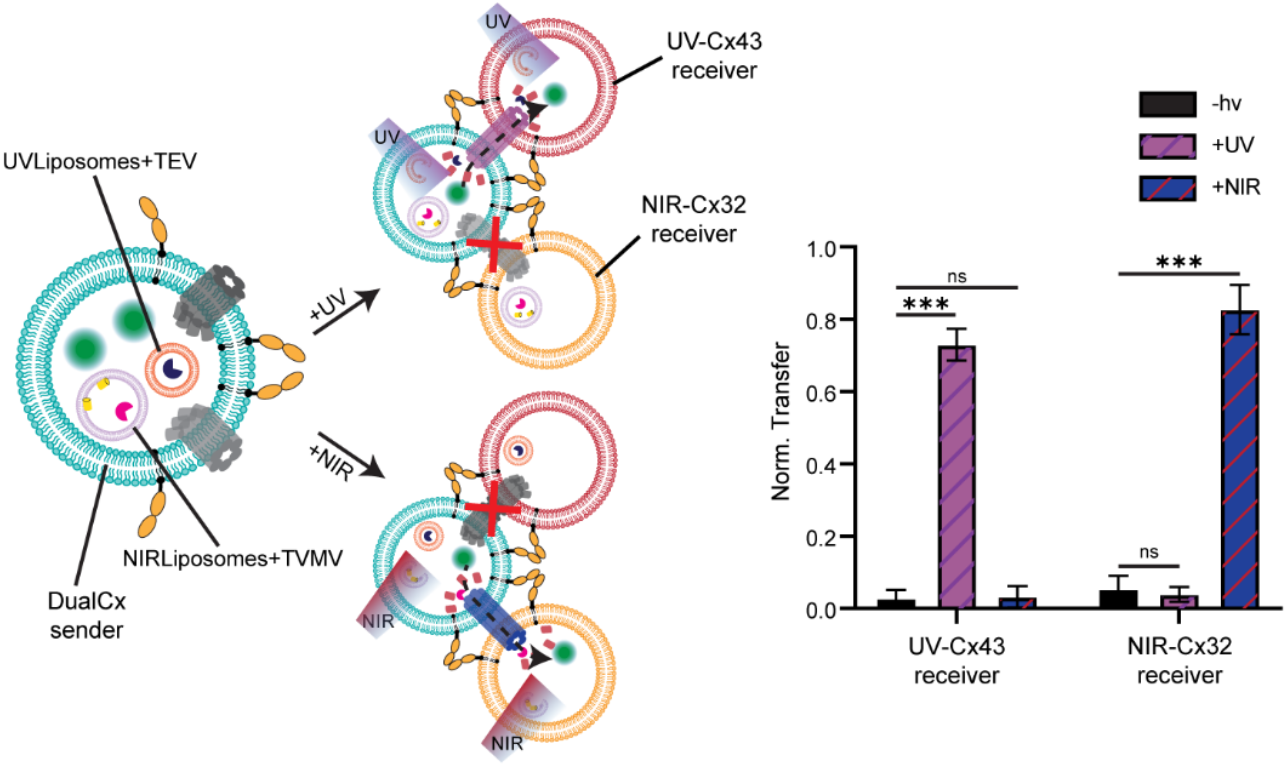
User-controlled communication network formation with a dual connexin-expressing SC. Scheme of DualCx sender SCs (left) that co-express non-functional TEVrecCx43 and TVMVrecCx32 connexons and co-encapsulate UV-responsive (UVLiposome+TEV) and NIR-repsonsive (NIRLiposome+TVMV) liposomes. DualCx senders are adhered with UV-Cx43 and NIR-Cx32 SCs that encapsulate UV-responsive or NIR-responsive, respectively. UV illumination specifically activates transfer between DualCx senders and UV-Cx43 receivers, while NIR illumination results in transfer to the NIR-Cx32 receivers. Normalized dye transfer to UV-Cx43 and NIR-Cx32 receivers under no, UV or NIR illumination (right). Error bars represent the s.d. of 3 independent trials, at least 30 receivers were analyzed per trial. Asterisks represent statistically significant differences (two-tailed unpaired t test, ***p < 0.001, n.s. p > 0.05).

Finally, we reasoned that our specific and selective transfer platform could be used to direct distinct reactants between SCs, ultimately leading to unique reaction products within a SC assembly (Fig. 6a) – a synthetic form of tissue signal transfer and processing but with user-defined control. To accomplish this, we encapsulated different fluorogenic azide reactants within SC lumens. In UV-Cx43 containing SCs, we included the fluorogenic azide CalFluor 647 (CF647), and in NIR-Cx32 containing SCs, we included the fluorogenic azide CalFluor 488 (CF488).^49^ Then, we generated DualCx SCs encapsulating a water soluble cyclooctyne, polyethylene glycol modified dibenzocyclooctyne (PEG-DBCO), that reacts with azides to form triazole products. We first adhered the three-SC population together using EC1-2 and verified that no reaction products formed since connexin channels remained in the inactive state, restricting the reactants to the separate SCs and leading to a dark state of the SC assembly (Fig 6b,c). Next, the SC assembly was illuminated with UV. Gratifyingly, we only observed formation of the fluorescent CF647 triazole product within the SCs, which stemmed from specific transfer from UV-Cx43 SCs to DualCx SCs and resulted in the red state of the SC assembly (Fig 6b,c). Similarly, after NIR illumination, we observed fluorescent CF488 triazole product within the SCs from specific transfer from NIR-Cx32 SCs to DualCx SCs, resulting in the green state of the system (Fig 6b,c). We also tested illuminating with both wavelengths simultaneously. In this case, we observed formation of both triazole products, which we describe as the yellow state (Fig 6b, c). In sum, our work describes an SC assembly that can specifically and selectively transfer different molecular species between individual SCs. By coupling transfer to reactive chemistry – a form of signal processing – we also showed that distinct SC assembly states can be formed, four in total, which is a critical step towards using SC communication networks for computing purposes beyond binary languages.

**Figure 6.**
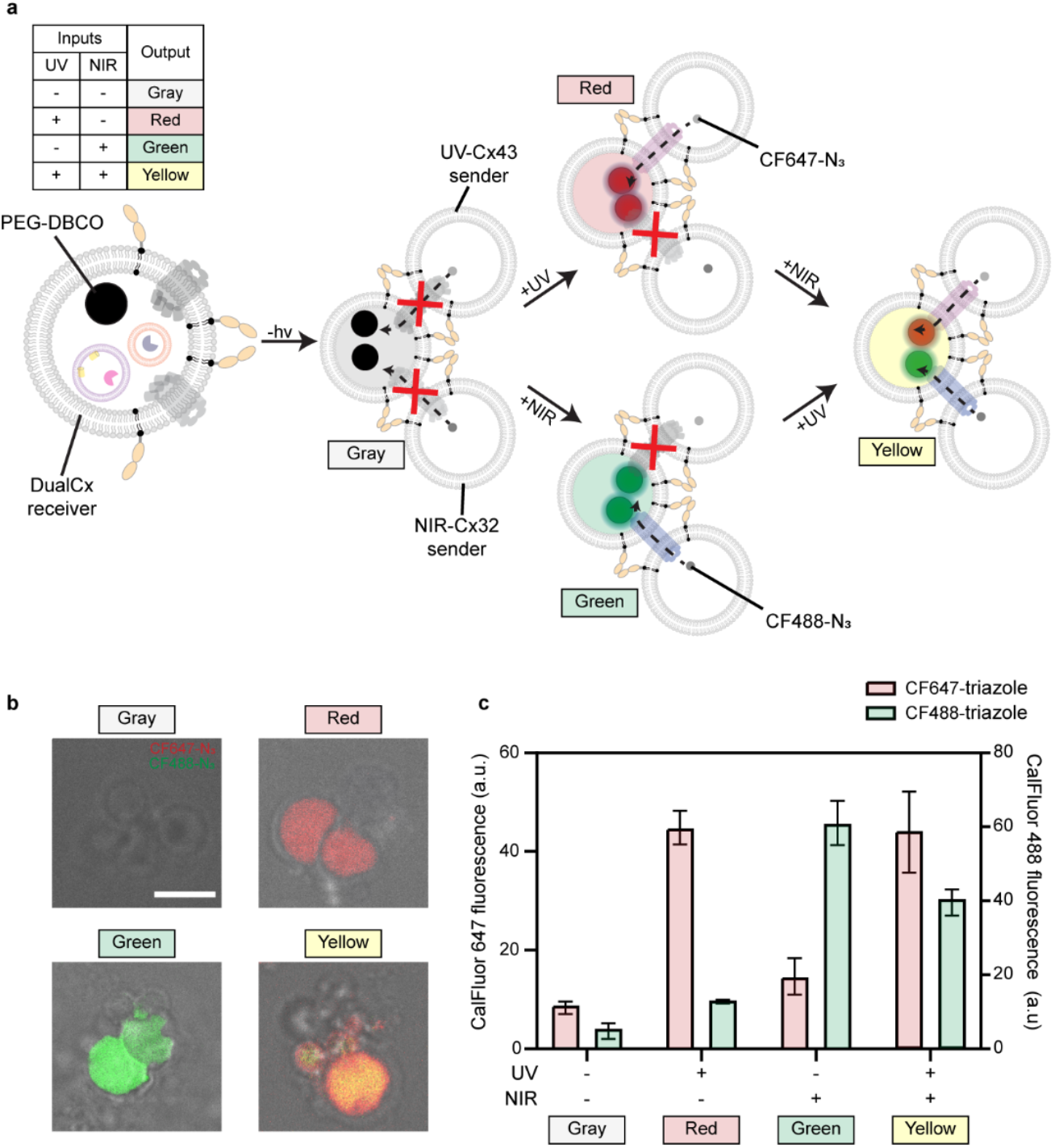
User-controlled production of different chemical outputs in dual connexin-expressing SCs. **a**, Scheme of DualCx receiver SCs that co-express the non-functional TEVrecCx43 and TVMVactiv-Cx32 connexons. These SCs encapsulate NIR- and UV-responsive liposomes along with polyethylene glycol-modified dibenzocyclooctyne (PEG-DBCO). These receivers are interfaced with UV-Cx43 or NIR-Cx32 sender SCs that are loaded with CalFluor 647 (CF647-N_3_)- and CalFluor488 (CF488-N_3_)-azide respectively. Using UV, activates Cx43-channels between UV-Cx43 senders and DualCx receivers, and leads to transfer of CF647-N_3_ to the receivers that produces a red signal upon a click reaction with PEG-DBCO. Conversely, NIR irradiation leads to green signal production due to the transfer of CF488-N_3_ across functional Cx32 channels between NIR-Cx32 senders and DualCx receivers. The truth table represents the relation between the light inputs and the output state that is produced in the receivers. **b**, Bright-field images merged with fluorescence channels show the different chemical states that the system achieves as a consequence of different input lights. **c**, Quantification of the amount of click-chemistry products (CF647- and CF488-triazoles) produced under different illumination states. Fluorescence of the respective chemical products is used as a proxy for their concentration. Error bars represent the s.d. of 3 independent trials, at least 35 receivers were analyzed per trial. Scale bar: 5 µm.

### Conclusion

Lipid bilayer vesicles offer notable chemical advantages as a material, such as long-term stability, compatibility with aqueous and biological environments, and differentiation between inner and outer environments. These properties position SCs, and other lipid bilayer structures, as ideal building blocks for communicative materials, as they can store molecular information at defined positions for long periods of time.^50^ Still, to transform SCs into a high-fidelity communication network demands a controllable conduit that spans two lipid bilayers, enabling the transfer of signals between SCs. This has been a long-sought, although largely unrealized, goal to date.^51–53^ Here, we provided one of the first examples of direct molecular transfer from one SC to another via connexin channel conduits, a key step toward constructing high-fidelity synthetic communication networks. Our approach relied on engineering gap junction connexins to undergo activation only in the presence of a light stimulus. We found that combining proteins from nature, such as connexins and E-cadherin, with non-natural systems, such as NIR-sensitive, gold nanorod-containing liposomes, endowed SCs with rapidly controllable capabilities, which would be hard to realize in either living cells or in purely non-natural materials. Further, by leveraging connexins that evolved to form incompatible channels, namely Cx43 and Cx32, orthogonal networks between SCs led to specific pathways for information transfer and were leveraged to produce unique reaction products within synthetic cells. Of note, we took advantage of orthogonal communication between SCs to create a SC logic system that produces four distinct outputs, i.e., reaction products, based on two inputs, namely wavelength of illumination. This could enable advanced logic operations beyond binary states using DualCx-based assemblies for future computing and circuit applications. As well, DualCx SCs represent a step towards an undifferentiated, pluripotent SC.^54^ As either UV or NIR light is applied to the material, DualCx SCs commit to one of two states irreversibly, analogous to stem cell differentiation, but determined entirely by the user.

Our platform provides researchers with a single starting SC material that can be patterned to any network architecture. Multiple co-existing signaling pathways would allow the integration of multi-stimuli in an initially inert material. With incompatible connexin isoforms, SC materials can be leveraged to form co-existing signaling pathways upon activation, ultimately giving rise to interpenetrating networks. Interestingly, activable SCs eliminate the need for low-throughput technologies that spatially pattern individual but different SCs in space, since illumination can generate the network architectures post-fabrication.^55^ As methods to prepare SCs and cell-free protein systems improve, we envision integrating additional natural or engineered connexin isoforms into single SCs to transfer many different signals that can be processed in subsequent SCs, for example via various reactive chemistries, for applications in regenerative medicine, sensing, and soft computing.

## Methods

### Materials and general methods

All chemical reagents were of analytical grade, obtained from commercial suppliers and used without further purification, unless otherwise noted. 4-(2-hydroxyethyl)-1-piperazineethanesulfonic acid (HEPES), Tris hydrochloride (Tris·HCl), sodium chloride (NaCl), calcium chloride (CaCl_2_), lanthanum chloride heptahydrate (LaCl_3_·7H_2_O), imidazole and OptiPrep Density Gradient Medium were purchased from Sigma-Aldrich. 1-palmitoyl-2-oleoyl-glycero-3-phosphocholine (POPC), 1,2-dioleoyl-sn-glycero-3-[(N-(5-amino-1-carboxypentyl)iminodiacetic acid)succinyl] (nickel salt) (DGS-NTA(Ni)), 1,2-dioleoyl-sn-glycero-3-phosphoethanolamine-N-(lissamine rhodamine B sulfonyl) (ammonium salt) (Liss Rhod PE), 1,2-dipalmitoyl-sn-glycero-3-phosphocholine (DPPC), 1,2-bis(10,12-tricosadiynoyl)-sn-glycero-3-phosphocholine (DC(8,9)PC), and 1,2-distearoyl-sn-glycero-3-phosphoethanolamine-N-[methoxy(polyethylene glycol)-2000] (DSPE-PEG(2000)) were obtained from Avanti Polar Lipids. 1,2-Dioleoyl-sn-glycero-3-phosphoethanolamine labeled with either Atto 390 (DOPE-Atto390) or Atto 647 (DOPE-Atto647) were obtained from ATTO-TEC GmbH. Gap26 peptide was purchased from Tocris Bioscience. Alexa Fluor 647-N-Hydroxysuccinimide (NHS) Ester and Alexa Fluor 488-NHS Ester were obtained from ThermoFisher Scientific. PURExpress *In Vitro* Protein Synthesis Kit and Tobacco Etch Virus (TEV) Protease were purchased from New England Biolabs. Recombinant C4 (TVMV) Protease was purchased from Abcam. Gold nanorods with citrate as a stabilizing agent with a ∼4.1 aspect ratio (length, ∼41 nm) was purchased from Nanopartz. Sensolyte 520 TEV Protease Assay Kit Fluorimetric was obtained from AnaSpec. Dibenzocyclooctyne conjugated with a polyethylene glycol linker (DBCO-PEG24) was purchased from BroadPharm. CalFluor 647- and CalFluor 488-azides were obtained from Vector Laboratories, Inc.

Fluorescence imaging was carried out on an Eclipse Ti2 microscope (Nikon) equipped with 405/488/560/640nm lasers (Andor), a Dragon 500 high speed spinning disk confocal module (Andor), and a Zyla 4.2 sCMOS camera (Andor). Fluorescence micrographs of vesicles were acquired with a 60x objective (Nikon, NA 1.49 TIRF) and were analyzed using Fiji (ImageJ). Brightness/contrast for all analyzed channels was kept consistent across images within a single experiment. Sample fluorescence was acquired with a BioTek Cytation 5 Imaging reader. For UV irradiation experiments, samples were placed in a 96-well plate (Nunc 96-well, Clear) at a distance of 6.5 cm from a UVP 3UV lamp (254 nm shortwave UV emission, 115V, 8W, Analytikjena). For near IR (NIR) experiments, the laser source used was a supercontinuum white-light laser source (SuperK Fianium FIU-15, NKT Photonics). An external RF driver was used to select the output wavelengths. An external RF driver was used to select the output wavelengths. Five channels of wavelengths (from 850 nm to 870 nm with increment of 5 nm) were turned on simultaneously. The output light was coupled by a single mode fiber and then collimated using an objective lens (Nikon OEM Microscope Objective Plan APO VC 20x Air 0.75 NA UV). The samples were placed in a 384-well plate (Greiner Microplate 384-well, Black) at a distance of 20 cm from the output collimated beam (12 mW/cm^2^).

Two-tailed unpaired t tests were performed using Prism 9 (GraphPad Software). For each experiment, the details of the replicates are noted in the figure legend. For all figures, ∗ p<0.05, ∗∗ p<0.01, ∗∗∗ p<0.001, ∗∗∗∗ p<0.0001, and n.s. p>0.05 were used.

### Plasmid construction

Cloning was performed using Gibson Assembly.^56^ Cx43-GFP and Cx32-GFP fusions were obtained by amplifying the Cx43^19^ and Cx32 genes (*rat* Cx32 gBlock, IDT) along with an eGFP gene using PCR and introducing them into the vector pRSET between XbaI and EcoRI restriction sites with Gibson Assembly (plasmids pRSET_Cx43-GFP and pRSET_Cx32-GFP). The protease-sensitive Cx43 and Cx32 variants were designed in a similar way by amplifying the Cx43-GFP and Cx32-GFP fusions along with an mCherry sequence using PCR and inserting them between the XbaI and EcoRI sites of the pRSET vector using Gibson Assembly (pRSET_mCherry-TEVrec-Cx43-GFP, pRSET_mCherry-TVMVrec-Cx32-GFP). Primers were used to introduce the TEV and TVMV recognition sites for the TEVrecCx43 and TVMVrecCx32 variants, respectively, and their sequences are listed in Supplementary Table 1.

Gene fragments encoding the first two (EC1-2; Asp1-Asp213) and three (EC1-3; Asp1-Phe332) extracellular domains of mouse E-cadherin were purchased as separate gBlocks from Integrated DNA Technologies (IDT). These fragments were then cloned into a pET28a vector containing a 6x His tag using Gibson Assembly to create the pET28a_EC1-2 and pET28a_EC1-3 plasmids.

### Protein expression and purification

Protein expression was adapted from Katsamba et al.^57^ Briefly, the EC1-2 and EC1-3 plasmids were transformed into *Escherichia coli* BL21 (DE3) and Rosetta II (DE3) pLysS cells and grown at 37 °C until an OD_600_ of 0.6 was reached. Expression was induced by using either 1 mM (for EC1-2) or 0.5 mM (for EC1-3) IPTG, and cells were subsequently grown for 16 h at 18 °C. Cells were then collected and resuspended in lysis buffer (10 mM Tris–HCl, pH 8.0, 500 mM NaCl, 3 mM CaCl_2_, 20 mM imidazole) containing protease inhibitors (ThermoScientific) and DNase I (ThermoScientific). Cells were lysed first using a Dounce homogenizer and then with sonication (Branson SFX250 Sonifier, 30% amplitude) for 6 min. Lysates were then clarified by centrifugation at 18,000 x *g* for 30 min at 4 °C and filtered through a 0.45-µm pore syringe filter. EC1-2 and EC1-3 were purified using affinity chromatography as follows. The clarified lysate was allowed to cycle over a HisTrap FF column (Cytiva) for 2 h at 4 °C using a BioLogic LP system (Biorad). The column was then washed with wash buffer containing 10 mM Tris–HCl, pH 8.0, 500 mM NaCl, 3 mM CaCl_2_, and 20 mM imidazole. The column was transferred to an Akta Pure 25 (Cytiva), and protein was eluted using a 20 CV gradient of 0-100% elution buffer (10 mM Tris–HCl, pH 8.0, 500 mM NaCl, 3 mM CaCl_2_, 500 mM imidazole). Eluted proteins were further purified by size exclusion chromatography using a Superdex 200 pg column (Cytiva) into buffer containing 20 mM Tris, pH 8.0, 150 mM NaCl, and 10 mM CaCl_2_. Recombinant Human E-Cadherin Protein (EC1-5) was purchased from R&D systems and reconstituted at 250 μg/mL in sterile 1X PBS (pH = 7.4, Gibco) according to the manufacturer’s instructions.

### Dye transfer in two-vesicle populations

Vesicles expressing connexin variants were prepared using a modified inverted emulsion process.^19^ Briefly, a 657 µM (total lipid) suspension of POPC:DGS-NTA(Ni):fluorescent lipid (94.9:5:0.1) in a silicone oil, mineral oil and decane (40:7:3) mixture was freshly prepared by sonication (30 min) before each experiment. The fluorescent lipid used for all sender vesicles was DOPE-Atto 390, whereas Liss Rhod-PE was used for receiver vesicles. 20 µL of the lipid-oil solution was added on top of 60 µL of the outer solution (25 mM HEPES, 100 mM NaCl, 2 mM CaCl_2_, 100 µM LaCl_3_, pH 7.4) in a 1.5 mL tube and allowed to rest for 30 min. 7 µL of the inner vesicle solution containing PURExpress system (solutions A and B), plasmid DNA (70 ng), 10 µM Alexa Fluor 647-COOH (AF647, added only in sender vesicles) and 10% OptiPrep was prepared at 4 °C in buffer (5 mM HEPES, 30 mM NaCl, pH = 7.4). 5 µL of the inner solution was added to 80 µL of the lipid-oil solution and vortexed for 50 s to create an emulsion. Next, the emulsion was gently layered over the lipid monolayer interface in the 1.5 mL tube. Centrifugation was immediately performed at 4 °C for 10 min at 2400 x *g*. After centrifugation, the tube was punctured at the bottom, and the vesicles in buffer were ejected into a fresh 1.5 mL tube. 40 µL of both the sender and the receiver vesicles were mixed in a fresh 1.5 mL tube through gentle pipetting. To this mixture, E-cadherin EC1-2 protein was added to a final concentration of 50 nM. Following a 10 min incubation on ice, the vesicle mixture was spun down at 4 °C for 10 min at 6000 x *g* to pellet both sender and receiver vesicles. The vesicle pellet was re-suspended in 30 µL of freshly prepared outer solution using a cut pipette tip and subsequently incubated at 37 °C for 2 h to facilitate connexin variant expression. Finally, the prepared vesicles were imaged using a 60x objective. The acquired fluorescence micrographs were analyzed in Fiji (ImageJ) to quantify the percentage of receiver vesicles exhibiting dye transfer. Only receiver vesicles in direct contact with sender vesicles were considered for analysis. Receiver vesicles were selected with the oval selection tool in Fiji. After calculating average receiver cell AF647 fluorescence, receiver cells with a lumenal dye fluorescence greater than 110% of the background fluorescence were classified as receivers with dye transfer. The above procedure applies to all dye transfer experiments unless otherwise noted.

For experiments with different E-cadherin variants, EC1-3 or EC1-5 were added to the vesicle outer solution (final conc. 50 nM) instead of EC1-2 in the above protocol. The above procedure was repeated by removing CaCl_2_ or adding 200 µM Gap26 peptide to the outer solution in order to verify whether dye transfer was calcium- or Cx43-dependent, respectively.

### Dye transfer in three-vesicle populations

Sender vesicles were prepared similar to the two-vesicle experiments with the exception that 10 µM Alexa Fluor 488-COOH (AF488) was encapsulated instead of AF647. The two receiver vesicle populations had membrane lipid compositions of POPC:DGS-NTA(Ni):Liss Rhod PE (94.9:5:0.1) and POPC:DGS-NTA(Ni):DOPE-Atto 647 (94.9:5:0.1), respectively. 40 µL of each sender and receiver vesicle solution was combined and gently mixed in a 1.5 mL tube using a pipette. To this mixture, EC1-2 was added to a final concentration of 50 nM and allowed to sit on ice for 10 min. Next, the vesicles were pelleted by centrifugation at 4 °C for 10 min at 6000 x *g*. The vesicle pellet was re-suspended in fresh 30 µL outer solution using a cut pipette tip, and subsequently incubated at 37 °C for 2 h to enable connexin expression. The sender vesicles expressed either Cx43 or Cx32 depending on the experiment (see Figure captions). The prepared vesicles were imaged using a 60x objective. The acquired images were manually analyzed in Fiji to calculate normalized dye transfer to receiver vesicles. Similar to above, we limited our analysis to receivers that were in direct contact with senders, classifying a lumenal dye fluorescence greater than 110% of the background as receivers that exhibited dye transfer. Normalization was performed with respect to dye transfer in two-vesicle population experiments. Percent transfer in two-vesicle populations with receivers lacking connexin plasmid was assigned a value of 0. A value of 1 was assigned to percent transfer in Cx43- or Cx32-expressing vesicle doublets.

### Preparation and characterization of light-sensitive liposomes

#### UV-responsive liposomes

A well-mixed solution of DPPC:DC(8,9)PC:DSPE-PEG2000 (79.5:20:0.5) lipids in chloroform (10 mg/mL) was placed under a stream of N_2_ gas to evaporate the chloroform. The film was kept in a vacuum desiccator for 2 h to remove residual chloroform from the lipid cake. Lipid films containing DC(8,9)PC were prepared in the absence of light and covered with foil. Next, the lipid film was hydrated with an aqueous solution (final lipid conc. 10 mg/mL) containing 100 Units of TEV protease, 1 mM DTT, 25 mM HEPES, 150 mM NaCl, pH 7.4, at 42 °C to form multilamellar vesicles. Multilamellar vesicles were then extruded 43 times through a 100 nm polycarbonate filter (Cytiva) at 42 °C to prepare liposomes encapsulating TEV protease, and the resulting liposomes were kept at 4 °C for 30 min. Next, the liposome samples were centrifuged at 16900 x *g* for 10 min at room temperature to separate any large lipid aggregates from liposomes. Finally, liposomes in the supernatant were purified using a micro spin G-50 column (Cytiva) to remove any unencapsulated TEV from the liposome solution.

#### NIR-responsive liposomes

A well-mixed solution of DPPC:DSPE-PEG2000 (99.5:0.5) lipids in chloroform (10 mg/mL) was placed under a stream of N_2_ gas to evaporate the chloroform. The lipid film was kept in a vacuum desiccator for 2 h to remove any residual chloroform from the lipid cake. Next, the lipid film was hydrated with an aqueous solution (final lipid conc. 10 mg/mL) containing either 100 Units of TEV or TVMV protease, 3 pmol gold nanorods, 25 mM HEPES, 150 mM NaCl, pH 7.4, at 42 °C to form multilamellar vesicles. Multilamellar vesicles were then extruded 43 times through a 200 nm polycarbonate filter (Cytiva) at 42 °C to prepare liposomes encapsulating either TEV or TVMV protease. The resulting liposomes were kept at 4 °C for 30 min. Next, the liposome samples were centrifuged at 16900 x *g* for 10 min at room temperature to separate any large lipid aggregates from liposomes. Finally, liposomes in the supernatant were purified using a micro spin G-50 column to remove any unencapsulated protease and gold nanorods from the liposome solution.

#### TEV release kinetics from NIR-responsive liposomes

30 µL solution of TEV-loaded NIR-responsive liposomes was added to a 384-well plate (Greiner Microplate 384-well, Black) and either illuminated with NIR light for different durations of time (1, 3, 8, 15 and 24 min) or kept in the dark at room temperature. As well, a sample with unencapsulated soluble TEV was prepared in a 25 mM HEPES, 150 mM NaCl, pH 7.4 buffer. Next, each of the illuminated and non-illuminated samples as well as the TEV-only sample were diluted 2-fold in 25 mM HEPES, 150 mM NaCl, pH 7.4 buffer. 50 µL of these diluted solutions was added to a 96-well plate (Corning 96-well, Black). 0.5 µL of TEV substrate (5-FAM-ENLYFQS-QXL 520 quencher, 100X) was diluted to 50 µL in buffer (25 mM HEPES, 150 mM NaCl, pH 7.4) and then added to each of the irradiated and non-irradiated samples. The mixtures were incubated at 30 °C in the plate reader, and 5-FAM fluorescence values were obtained every 5 min for a total of 15 min (λ_ex/em_ = 492/518 nm). The percentage of TEV release for the NIR and for the non-NIR irradiated liposomes was calculated by comparing the fluorescence value increase to that of unencapsulated soluble TEV in buffer.

#### Dynamic Light Scattering

Liposome diameter was determined via dynamic light scattering using a Zetasizer Nano ZS size analyzer (Malvern Panalytical). 50 µL of liposome solution was diluted with 950 µL of 25 mM HEPES, 150 mM NaCl, pH 7.4 buffer, and size distributions of particles were measured via three runs with 16 measurements per run.

#### Light-activated dye transfer in two-vesicle populations

#### UV-activated dye transfer

1 µL of UV-responsive liposomes along with the PURExpress system, TEVrec-Cx43 plasmid (70 ng), 10% OptiPrep and 10 µM AF647 in buffer (5 mM HEPES, 30 mM NaCl, pH 7.4) were mixed to form 7 µL of the inner solution of the sender vesicles (UV-Cx43 sender). Receiver vesicles (UV-Cx43 receiver) had a similar inner solution composition but lacked the AF647 dye. The sender and receiver vesicles were adhered using EC1-2 (50 nM) and subjected to protein expression at 37 °C for 2 h. Following TEVrecCx43 expression, the two-vesicle population was illuminated with 254 nm UV light for 10 min. Sender and receiver vesicles prepared similarly but i) encapsulating 5 Units of soluble TEV, ii) not exposed to UV light, or iii) not encapsulating UV-responsive liposomes served as control samples. Vesicles were imaged using a 60x objective and subsequently analyzed for dye transfer using Fiji. Normalization was performed with respect to dye transfer in two-vesicle populations with (1) and without (0) Cx43 expression.

#### NIR-activated dye transfer

The experimental procedure was similar to that of UV-activated transfer, with the primary difference being the use of 1 µL of NIR-responsive liposomes for the vesicle inner solution and NIR light source for activation. Following TEVrecCx43 expression, the two-vesicle population was illuminated with NIR light for 15 min. Sender and receiver vesicles prepared similarly but i) encapsulating 5 Units of soluble TEV, ii) not exposed to NIR light, or iii) not encapsulating NIR-responsive liposomes served as control samples. All vesicles were imaged using a 60x objective and subsequently analyzed for dye transfer using Fiji. Normalization was performed with respect to dye transfer in two-vesicle populations with (1) and without (0) Cx43 expression. For NIR-activated transfer experiments with TVMVrecCx32, 1 µL of TVMV-loaded NIR-responsive liposomes were encapsulated in the sender (NIR-Cx32 sender) and receiver (NIR-Cx32 receiver) vesicles instead. Normalized transfer was calculated with respect to dye transfer in two-vesicle populations with (1) and without (0) Cx32 expression.

### Orthogonal light-activated dye transfer in three-vesicle populations

0.5 µL each of UV- and NIR-responsive liposomes along with the PURExpress system, TEVrecCx43 plasmid (70 ng), TVMVrecCx32 plasmid (70 ng), 10% OptiPrep and 10 µM AF488 in buffer (5 mM HEPES, 30 mM NaCl, pH 7.4) were mixed to form 7 µL of the inner solution of the DualCx sender vesicles. The two receiver vesicle populations comprised of UV-Cx3 receivers and NIR-Cx32 receivers. Sender vesicles were adhered with both receiver vesicles using EC1-2 (50 nM). Following protein expression at 37 °C for 2 h, the three-vesicle population was illuminated with either UV for 10 min or NIR for 15 min. Finally, vesicles were imaged using a 60x objective, and receiver vesicles were analyzed for dye transfer using Fiji as described previously (see Dye transfer in three-vesicle populations). Dye transfer was normalized with respect to percent transfer from DualCx senders to either Cx43- or Cx32-receiver vesicles (see Supplementary Fig. 11) in the presence of either TEV or TMVM protease (1). Similarly, dye transfer from DualCx senders to either Cx43- or Cx32-receivers in the absence of any protease served as the lower bound (0) for normalization.

### Orthogonal light-activated fluorogenic click-reactions in three-vesicle populations

0.5 µL each of UV- and NIR-responsive liposomes along with the PURExpress system, TEVrecCx43 plasmid (70 ng), TVMVrecCx32 plasmid (70 ng), 10% OptiPrep and 200 µM PEG-DBCO were mixed to form 7 µL of the inner solution of the DualCx receiver vesicles. UV-Cx43 senders loaded with 10 µM CalFluor 647-azide, and NIR-Cx32 senders encapsulating 10 µM CalFluor 488-azide constituted the two sender vesicle populations. Receiver vesicles were adhered with both sender vesicles using EC1-2 (50 nM). Following protein expression at 37 °C for 2 h, the three-vesicle population was illuminated with either UV for 10 min or NIR for 15 min. Finally, vesicles were imaged using a 60x objective, and receiver vesicles were analyzed for CalFluor transfer and reaction with PEG-DBCO using Fiji. Receiver vesicles with either CalFluor 647 or CalFluor 488 signal were selected with the oval selection tool in Fiji. The mean CalFluor 647 and CalFluor 488 fluorescence in these receivers was calculated and plotted for each illumination condition.

## Acknowledgements

This work was supported by grants from the National Institutes of Health (R35GM142941 to B.B. and R35GM139531 to J.C.S.), the National Science Foundation (Grant 2218467 to B.B. and J.C.S.), the Welch Foundation (Grant F-2055-20240404 to B.B.), the David and Lucile Packard Foundation (2023-76153 to B.B.), the Center for Dynamics and Control of Materials: an NSF MRSEC (Grant 2308817 to B.B. and J.C.S.), and the Chan Zuckerberg Initiative (Grants 2021-225666 and 2023-321173 to S.C.). A.J.L. is supported by an NSF predoctoral fellowship.

