## Supplementary Information for "Light-Controlled Synthetic Communication Networks via Paired Connexon Nanopores"

**Supplementary Figures**

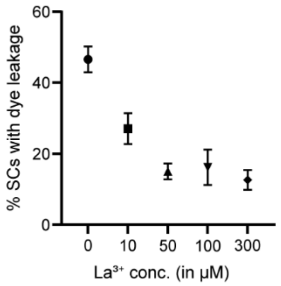


**Fig. S1.** Quantification of dye leakage above background leakage from Cx43-expressing SCs upon addition of different $La^{3+}$ concentrations to the SC outer solution. Background leakage refers to dye leakage from SCs lacking Cx43 expression. Error bars represent the s.d. of 3 independent trials, at least 30 SCs were analyzed per trial.

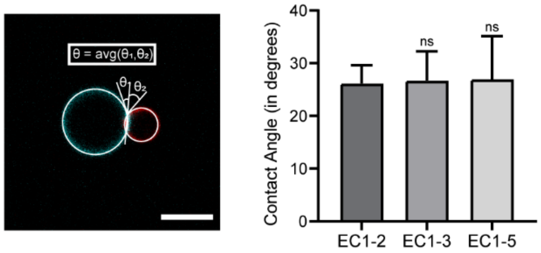


**Fig. S2.** Contact angle (θ) measurements of Cx43-expressing SC doublets adhered using different E-cadherin variants. Similar contact angles across the different variants indicate comparable binding energy. Error bars represent the s.d. of 3 independent trials, at least 15 SC doublets were analyzed per trial. Asterisks represent statistically significant differences (two-tailed unpaired t test, n.s. $p > 0.05$). Scale bar: 5 µm.

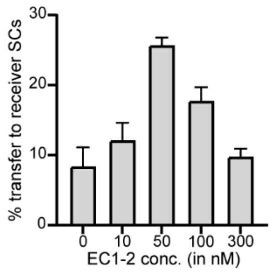


**Fig. S3.** Quantification of dye transfer above background transfer across Cx43 channels upon addition of different EC1-2 concentrations to the SC outer solution. Background here refers to dye transfer between senders and receivers lacking Cx43 expression. Error bars represent the s.d. of 3 independent trials, at least 30 SCs were analyzed per trial.

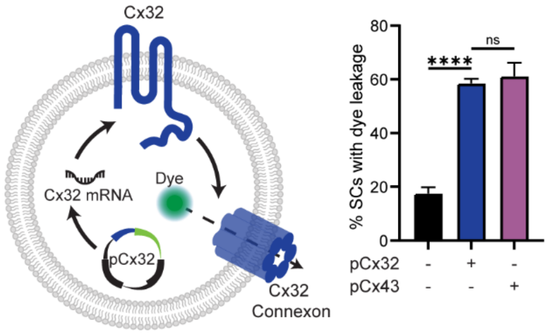


**Fig. S4.** Scheme of dye leakage from functional Cx32 connexons in SCs (left). Quantification of dye release from Cx32-GFP (right) expressing SCs reveals comparable connexon activity as Cx43-GFP expressing SCs. Error bars represent the s.d. of 3 independent trials, at least 35 SCs were analyzed per trial. Asterisks represent statistically significant differences (two-tailed unpaired t test, ****$p < 0.0001$, n.s. $p > 0.05$).

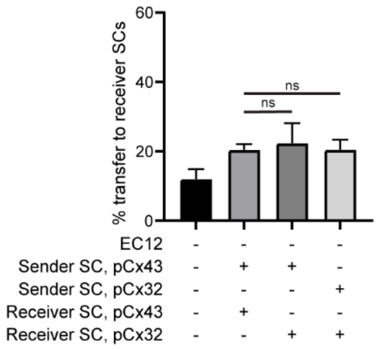


**Fig. S5.** Quantification of dye transfer between Cx43- or Cx32-expressing sender and receiver SCs in the absence of EC1-2 confirm the significance of adhesion in efficient inter-SC dye transfer. Error bars represent the s.d. of 3 independent trials, at least 45 receivers were analyzed per trial. Asterisks represent statistically significant differences (two-tailed unpaired t test, ns > 0.05).

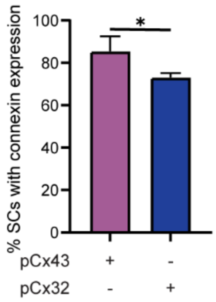


**Fig. S6.** Quantification of SCs expressing either Cx43-GFP or Cx32-GFP point to a lower Cx32 expression compared to Cx43 in SCs. Error bars represent the s.d. of 3 independent trials, at least 35 SCs were analyzed per trial. Asterisks represent statistically significant differences (two-tailed unpaired t test, * $p > 0.05$).

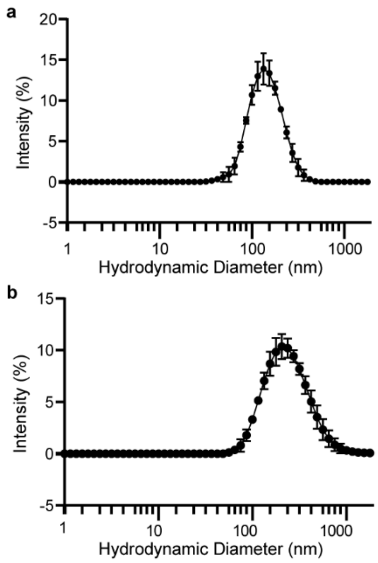


**Fig. S7.** Characterization of **a,** UV-responsive and **b,** NIR-responsive liposomes by Dynamic Light Scattering. Error bars represent s.e.m. (n = 3).

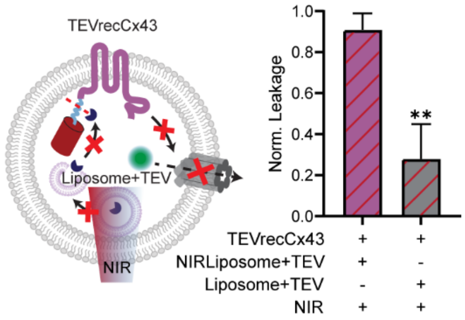


**Fig. S8.** Scheme showing TEVrecCx43 inactivity despite NIR illumination (left). TEV-loaded liposomes (Liposome+TEV) do not include gold nanorods, thus rendering them unresponsive to NIR. As a result, no TEV is released and the connexon remains non-functional. Normalized dye leakage upon NIR illumination in the presence and absence of liposomes functionalized with gold nanorods (right). Normalization was performed with respect to dye leakage from SCs expressing (1) or lacking (0) Cx43. Error bars represent the s.d. of 3 independent trials, at least 50 SCs were analyzed per trial. Asterisks represent statistically significant differences (two-tailed unpaired t test, **p < 0.01).

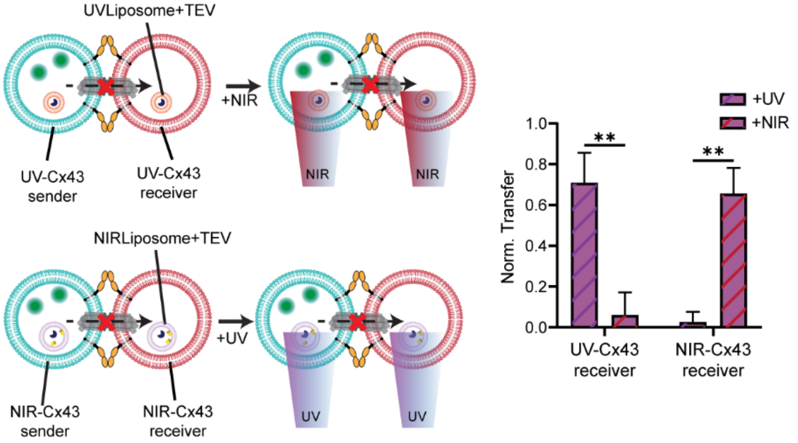


**Fig. S9.** Scheme of adhered TEVrecCx43-expressing sender and receiver SCs with incompatible activation wavelength. NIR does not activate UV-responsive liposomes, and similarly NIR-responsive liposomes are unresponsive to UV (left). Normalized dye transfer between TEVrecCx43 expressing senders and receivers that encapsulate either UV- or NIR-responsive liposomes upon UV or NIR illumination (right). Normalization was done with respect to dye transfer between senders and receivers expressing (1) or lacking (0) Cx43. Error bars represent the s.d. of 3 independent trials, at least 30 receivers were analyzed per trial. Asterisks represent statistically significant differences (two-tailed unpaired t test, $^{**}p < 0.01$).

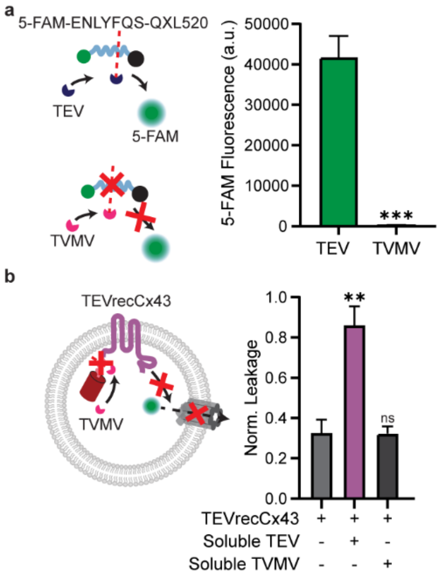


**Fig. S10. a,** Scheme of TEV protease cleavage of substrate sequence, which cannot be processed by TVMV (left). TEV substrate, labeled with a fluorescent dye, 5-FAM, and a quencher, QXL520, were added to either a TEV or TVMV-containing solution. Cleavage of the TEV substrate leads to production of 5-FAM signal. Comparison of 5-FAM fluorescence in the presence of either TEV or TVMV protease (right). Error bars represent the s.d. of 3 independent trials. **b,** Scheme of inactive TEVrecCx43 connexons in the presence of soluble TVMV (left). TVMV is unable to cleave the mCherry domain of TEVrecCx43, and as a result the connexon nanopore remains non-functional. Normalized dye leakage from TEVrecCx43 expressing SCs that encapsulate either TEV or TVMV protease (right). Normalization was performed with respect to leakage from SCs expressing (1) or lacking (0) Cx43. Error bars represent the s.d. of 3 independent trials, at least 30 SCs were analyzed per trial. Asterisks represent statistically significant differences (two-tailed unpaired t test, $^{**}p < 0.01$, $^{***}p < 0.001$).

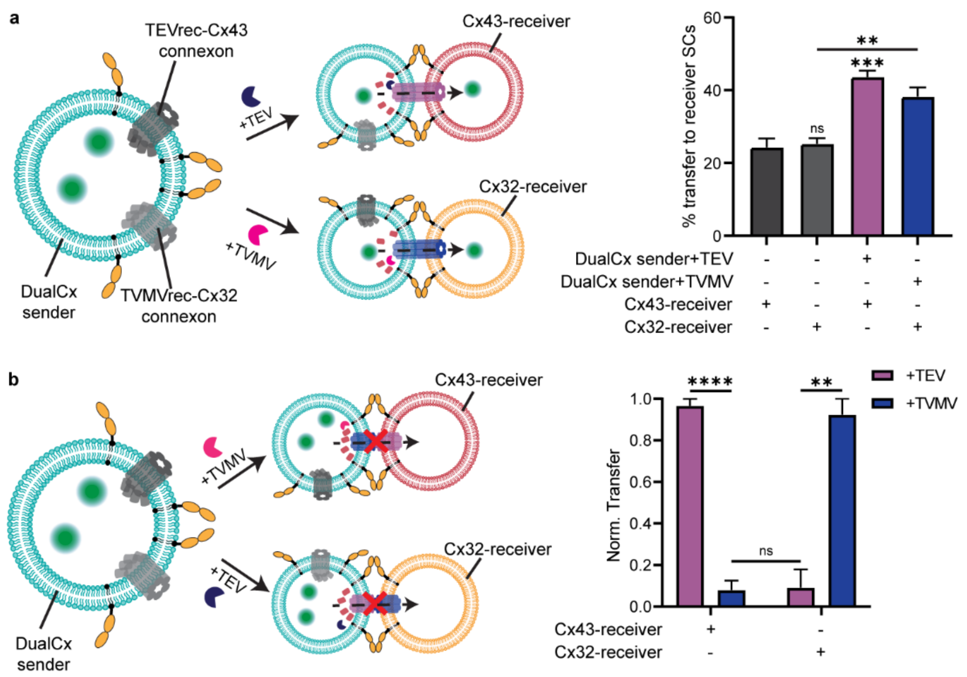


**Fig. S11. a,** Scheme of DualCx sender SCs co-expressing non-functional TEVrecCx43 and TVMVrecCx32 (left). Upon the addition of the appropriate protease to their lumen, DualCx sender can communicate specifically with either Cx43- or Cx32- expressing receiver SCs. Quantification of dye transfer to Cx43- or Cx32- expressing receivers in the absence and presence of TEV or TVMV protease (right). Error bars represent the s.d. of 3 independent trials, at least 30 receivers were analyzed per trial. **b,** Scheme of DualCx senders and Cx43- or Cx32-receivers with mismatched connexons (left), which would lead to a lack of channel formation. TEV activates TEVrecCx43 whereas TVMV activates TVMVrecCx32. Minimal dye transfer is observed when the former is interfaced with Cx32-receivers and the latter with Cx43-receivers. Normalized dye transfer to Cx43- or Cx32-expressing receivers from DualCx sender in the presence of either TEV or TVMV protease. For DualCx senders encapsulating TEV, normalization was performed with respect to dye transfer between senders and receivers expressing (1) or lacking (0) Cx43. For DualCx senders loaded with TVMV, normalization was performed with respect to dye transfer between senders and receivers expressing (1) or lacking (0) Cx32. Error bars represent the s.d. of 3 independent trials, at least 30 receivers were analyzed per trial. Asterisks represent statistically significant differences (two-tailed unpaired t test, $^{**}p < 0.01$, $^{***}p < 0.001$, $^{****}p < 0.0001$, n.s. $p > 0.05$).

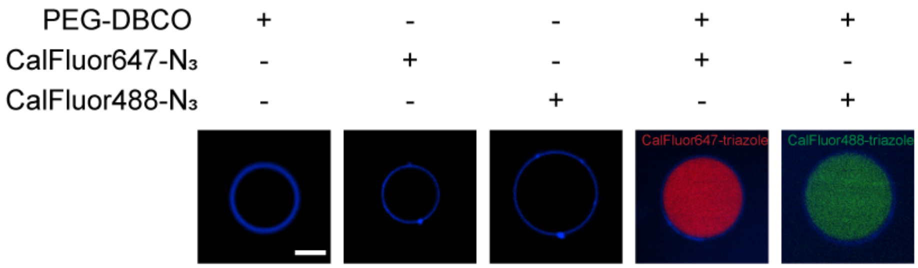


**Fig. S12.** Fluorescence micrographs showing that the Cu-free click reaction between polyethylene glycol modified-dibenzocyclooctyne (PEG-DBCO) and CalFluor-azides is compatible with lipid vesicles (blue) used in this study. Further, this reaction produces fluorescent products only when both the reactants are present inside a vesicle. Scale bar: 5 µm.

**Supplementary Table**

| Cx variant | Oligonucleotide sequence (5'-3') |
|---|---|
| TEVrecCx43 | Forward:<br>AGAAAACCTGTATTTTCAGAGCATGGGTGACTGGAGTGCC<br>Reverse:<br>TCACCCATGCTCTGAAAATACAGGTTTTCTTTGTATAATTCGTCCATTCCACCTG |
| TVMVrecCx32 | Forward:<br>AGAGACTGTTCGTTTCCAATCGATGAACTGGACTGGACTTTATACCC<br>Reverse:<br>CAGTTCATCGATTGGAAACGAACAGTCTCTTTGTATAATTCGTCCATTCCACCTG |

**Table 1.** Primer sequences used to introduce the TEV and TVMV recognition sites for the TEVrecCx43 and TVMVrecCx32 variants. Highlighted bases represent hybridization sites for PCR.
